# Co-expression of Bovine leukemia virus and Bovine foamy virus-derived miRNAs in naturally infected cattle

**DOI:** 10.1101/2025.02.21.639540

**Authors:** MI Petersen, G Suarez Archilla, CV Perez, DD Gonzalez, KG Trono, C Mongini, JP Jaworski, HA Carignano

## Abstract

Among the viruses that encode miRNAs and infect cattle such as Bovine leukemia virus (BLV), Bovine foamy virus (BFV) and Bovine herpesviruses (BoHV), BLV has gained attention due to the critical role that BLV-miRNAs may play in inducing lymphosarcoma in infected animals. BLV is highly prevalent in the Americas and negatively affects dairy herds, primarily due to restrictions on the commercialization of dairy products from infected animals and a decrease in milk production. Mixed infections involving BLV and BFV appear to be common in cattle. Considering the ability of foamy viruses to cross species barriers, preventing their presence within the food chain is essential. We identified the co-expression of seven BLV-derived miRNAs (blv-miR-B1-3p, blv-miR-B2-5p, blv-miR-B2-3p, blv-miR-B3-5p, blv-miR-B3-3p, blv-miR-B4-3p and blv-miR-B5-5p) and three BFV-derived (bfv-miR-BF1-5p, bfv-miR-BF1-3p, and bfv-miR-BF2-5p) in naturally BLV-infected cows. Besides, seven differentially expressed bovine miRNAs (bta-miR-375, bta-miR-133a, bta-miR-677, bta-miR-1, bta-miR-3613a, bta-miR-9-5p and bta-miR-95) were identified between cows with high BLV proviral load and uninfected counterparts (fold change > |1.5| and q-value < 0.05). A comprehensive functional analysis of protein-protein interaction networks for genes targeted by both viral and host-derived miRNAs highlighted key pathways implicated in tumorigenesis and immune response. Although BLV and BFV derived miRNAs target different genes, their functional convergence may reflect coordinated viral modulation of host cellular processes, raising important concerns regarding their potential influence on disease severity and the increased dissemination of BFV. These findings offer new perspectives for creating diagnostic and treatment approaches to manage viral persistence and tumorigenesis in cattle.

**Importance:** The Bovine leukemia virus (BLV) and Bovine foamy virus (BFV) are retroviruses that encode miRNAs and infect cattle. While the role of BFV-derived miRNAs remains unclear, BLV-miRNAs have gained attention for their potential involvement in oncogenesis. Mixed BLV-BFV infections are common and given foamy viruses’ potential to cross species barriers, it is essential to prevent their presence in the food chain. We reported the co-expression of three BFV-derived miRNAs and seven BLV-derived miRNAs in naturally infected cattle. A functional analysis of the protein-protein interaction graph of genes potentially targeted by both viral and host-derived miRNAs revealed key metabolic pathways associated with tumorigenesis and immune response regulation. The co-expression of BLV and BFV miRNAs suggests a potential functional interactions that may influence disease progression and BFV dissemination. These findings offer opportunities for developing diagnostic and therapeutic strategies to control viral persistence and tumor development in cattle.

## Introduction

The study of virus-host interactions has traditionally focused on how viral proteins modulate host proteins to hijack cellular processes in favor of viral replication and pathogenesis (1) However, like their host cells, many viruses encode regulatory non-coding RNAs, including small RNAs such as microRNAs (miRNAs), which provide new capacities for regulating host gene expression to ensure successful viral infection (2; 3; 4; 5). miRNAs are enzymatically processed small RNAs, typically 19–24 nucleotides (nt) long, that fine-tune gene expression at the post-transcriptional level (6). The regulatory function of miRNAs is primarily mediated by the RNA-induced silencing complex (RISC), which facilitates base-pairing between the mature miRNÁs 2–7 nucleotide seed region and the 3’ untranslated region (3’UTR) of the target messenger RNA (mRNA) (7; 8). The miRNA-mRNA interaction induces mRNA degradation by cellular endonucleases or translational repression (8; 9). It is estimated that miRNAs may regulate at least 60% of the human coding genome (10; 11).

Host-derived microRNAs (miRNAs) regulate viral infections by exhibiting a dual nature that can benefit either the virus or the host. For example, host miRNAs that directly target viral RNA often enhance the viral life cycle. Conversely, host miRNAs can exert an indirect negative effect on the virus by targeting mRNAs that encode essential host factors necessary for the viral cycle or for establishing immune responses and defense mechanisms (5; 12). Given their high versatility in fine-tuning gene expression and their small coding size, it is not surprising that viruses have evolved the ability to express their own miRNAs (13). Since the first report of miRNAs encoded by the Epstein-Barr virus (EBV) (14), miRNAs encoded by DNA viruses have proven to be the most abundant, particularly those expressed by the

Herpesviridae family. In contrast, RNA viruses that express miRNAs are far less common and are typically restricted to those with a DNA intermediate. Additionally, several miRNA-like molecules have been identified in both positive and negative-sense RNA viruses (3; 15; 16; 17; 18; 19; 20; 21; 22; 23; 24; 25; 26; 27; 28).

Usually, viral miRNAs follow the same biogenesis and effector pathways as host miRNAs. However, certain retroviruses, such as members of the Spumaretrovirinae subfamily (e.g., Simian foamy virus (SFV) and Bovine foamy virus (BFV)) and the Orthoretrovirinae subfamily (e.g., Bovine leukemia virus (BLV)), decouples miRNA transcription from genomic transcription by employing RNA polymerase III instead of RNA polymerase II (21; 22; 23). This mechanism likely prevents the auto-degradation of viral genomes mediated by endonucleases during miRNA biogenesis. (29; 30).

Generally, the functional role of viral miRNAs involves the downregulation of both viral and host proteins; however, upregulation would also occur—either indirectly through the modulation of gene that act as transcriptional repressors, or directly, as some miRNAs can function as transcriptional activators by binding to specific DNA sequences or interacting with factors that enhance gene expression (18; 31; 32). Collectively, these actions modulate gene expression to regulate the viral life cycle (e.g., promoting or inhibiting cell proliferation and apoptosis), evade the host immune response, and establish latent or persistent infections (4; 16; 33; 34; 35). This is particularly evident in retroviruses and herpesviruses, where viral protein expression is strongly restricted during latency. The production of a low-antigenicity regulatory molecule provides a significant advantage for viral persistence while evading host immune detection (13; 36).

Viruses that encode miRNAs and infect cattle include BLV (21), BFV (23), Bovine herpesvirus 1 (BoHV-1) (37), and Bovine herpesvirus 5 (BoHV-5) (38). BoHV-1 encodes at least ten miRNAs, two of which have been implicated in maintaining latency both *in vivo and in vitro* (39; 40). In contrast, the functional roles and target genes of the three BFV-derived miRNAs remain poorly understood (41). Among BLV-encoded miRNAs, blv-miR-B4-3p is the most extensevely studied. It shares a seed sequence with miR-29a, an oncomiR that targets tumor suppressors such as peroxidasin homolog (*PXDN*) and HMG-box transcription factor 1 (*HBP1*), as reported in murine tumor-induction studies (42; 43). *In vitro* reporter assays have demonstrated that blv-miR-B4-3p exerts a repressive effect on the expression of *HBP1* and *PXDN*. However, a negative correlation between *blv-miR-B4-3p* and *HBP1* expression has been documented, while levels of *PXDN* remained unchanged in ovine primary tumor B cells (21; 44). Additionally, recent findings indicate that *PXDN* expression is significantly downregulated in naturally BLV-infected cattle expressing blv-miR-B4-3p when compared to their non-infected counterparts. Notably, miR-29a expression remains unaffected in both groups (45).

The BLV prevalence reaches up to 85% in North America at farms-level and over 80% in lactating cows of Argentina’s main dairy region (46; 47; 48; 49). International trade restrictions of livestock products from affected herds impact negatively the economy (50), but greater losses stem from reduced milk production and earlier culling of asymptomatic BLV carriers compared to BLV-free herds (51; 52).

Mixed infections involving BLV and BFV appear to be common in cattle (53). Given the ability of foamy viruses (FVs) to cross species barriers and cause zoonotic infections in humans, their presence in the food chain may be a potential risk to both human and animal health. Furthermore, cattle infected with BLV are more susceptible to co-infections with BoHV-1 (54; 55; 56).

This study identified the expression of three BFV-derived miRNAs and seven BLV-derived miRNAs in naturally BLV-infected cattle through small RNA sequencing analysis. No miRNAs from BoHV-1/5 were found. Seven bovine miRNAs showed differential expression between cattle infected with BLV/BFV and those uninfected with BLV. A functional analysis of the protein-protein interaction network for genes potentially targeted by both viral and host-derived miRNAs revealed key metabolic pathways associated with tumorigenesis and immune response regulation.

## Methods

### Selection of Animals

BLV-infected and non-infected samples were obtained from a previously phenotyped population as described in Petersen et al. (2021) (57). Briefly, 129 adult Holstein cows (over 3 years old sharing the same lactation period) from a dairy farm in the central region of Argentina -where the average individual prevalence of BLV exceeds 80% (49)-were initially screened. Since antibody levels have been reported to reflect proviral load (PVL) *in vivo* (49; 58), anti-BLV ELISA (see below) was assesed at −10 months (T1) and −5 months (T2) before final sampling; mean percentage of reactivity (PR) was 122.7 ± 34.8 at T1 and 146.3 ± 55.6 at T2. Animals in the highest PR quartile (Q) (Q4: T1 = 148.6–178.6%, T2 = 194.4– 239.6%) and those testing negative at both times were selected for further proviral load (PVL) quantification via qPCR (details provided later) at −3 months (T3) and at the time of sampling (T4). Based on the consistency of PVL results, four cows with persistently high PVL (HPVL) and three cows with consistently negative qPCR results were ultimately selected for small RNA sequencing. All sample collection and handling procedures adhere to the recommendations and guidelines of the Institutional Committee for the Care and Use of Experimental Animals at the Instituto Nacional de Tecnología Agropecuaria.

### Isolation of Peripheral Blood Mononuclear Cells (PBMCs)

Fresh blood samples from animals were collected via jugular venipuncture and supplemented with EDTA (225 µM). Peripheral blood mononuclear cells (PBMCs) were isolated on the same day of collection using Ficoll-Paque Plus (GE Healthcare, Uppsala, Sweden) density gradient centrifugation, following the manufacturer’s protocol. The plasma fraction was kept for anti-BLV ELISA serology. After isolation, PBMCs were preserved in RNAlater solution (Ambion, Austin, TX) and stored at −80°C until further use.

### Serology

The anti-BLV ELISA assay, as described by Trono et al. (2001) (59), was employed to identify BLV-infected animals. The whole BLV viral particle served as the antigen for detecting plasma anti-BLV antibodies in each sample. Briefly, a sample-to-positive (S/P) ratio, referred to as the percentage of reactivity (PR), was calculated using the following formula: S/P = [(OD_Sample_ - OD_WS-)_ / (OD_WS+1/7_ -OD_WS-)]_ × 100, where OD refers to optical density, WS-indicates the negative serum, and WS+1/7 is the international standard weak positive control serum (diluted 1:7 in negative serum). Samples with a PR greater than 25% were considered positive (BLV(+)), while those below were regarded as negative (BLV(-)).

### BLV proviral load (PVL) Quantification

Genomic DNA was extracted from PBMCs using the High Pure PCR Template Preparation Kit (Roche, Penzberg, Germany) following the manufacturer’s instructions. The quality and concentration of genomic DNA from whole blood samples, extracted with the Blood Genomic DNA AxyPrep™ kit (Axygen Biosciences, Union City, USA), were assessed using a microvolume spectrophotometer (NanoDrop™ Technologies, Inc., Wilmington, USA).

A BLV POL gene-based PVL qPCR assay based on the SYBR Green dye detection system was conducted as described by Petersen et al. (2018) (60). Each 25 µL qPCR reaction contained Fast Start Universal SYBR Green Master Mix (2×; Roche), forward and reverse primers (800 nM; BLVpol_5f:5′-CCTCAATTCCCTTTAAACTA-3′ and BLVpol_3r:5′-GTACCGGGAAGACTGGATTA-3′; Thermo Fisher Scientific), and 200 ng of genomic DNA template. Amplification and detection were carried out using a Step One Plus system (Applied Biosystems, Foster City, CA). The specificity of each BLV-positive reaction was confirmed by melting temperature dissociation curve (Tm) analysis. Based on the assumption of a low natural infection rate (1% of BLV-infected cells in the peripheral blood), PVL values below 1,500 copies/µg of total DNA were classified as low, while values above this threshold were classified as high (61).

### Small RNA Sequencing

Total RNA was extracted from PBMCs using the miRNeasy kit (Qiagen, Hilden, Germany) according to the manufacturer’s protocol. RNA quality (integrity) and concentration were assessed via digital electrophoresis on an Agilent 2200 TapeStation system (Agilent Technologies, Santa Clara, US).

The QIAseq miRNA Library Kit (Qiagen, Hilden, Germany) was used for miRNA sequencing library construction, which integrates Unique Molecular Indices (UMIs) tags into the adapters during the cDNA synthesis step. These UMI tags help to mitigate PCR amplification biases and sequencing artifacts by allowing identification and collapsing reads with the same amplification origin. A specific 8-nt barcode was assigned to each sample for multiplexing. The multiplexed libraries were pooled to an equal molar concentration and processed in a NovaSeq run (Illumina, San Diego, US) with a 75x50 bp configuration (300–400 million reads pairs per lane).

### Small RNA sequencing reads processing and deduplication

The quality control of sequencing reads was performed sequentially to ensure reliable reads for unbiased miRNome identification and quantification. The 12-nt UMI pattern was extracted from each read using the umi_tools v1.1.6 software (62). Forward (Fw) and reverse (Rv) reads were processed independently before collapsing. Reads with low quality (quality cutoff < 30), a minimum length < 16 nt, the presence of ambiguous bases (N) or adapter sequences were removed using Cutadapt v5.0 (63).

Singleton reads resulting were discarded, and only paired Fw and Rv reads were considered. Forward read sequences were aligned to the Rv-complement and merged into a single consensus read using BBMap v25.85 (64) with a minimum insert size of 16 nt and a minimum overlap of 18 nt. After merging, Qiagen-specific adapter sequences were removed using Cutadapt v5.0 (63).

A stringent mapping strategy was employed for read deduplication using Bowtie2 v2.5.4 (65). Reads were aligned to the bovine reference genome ARS-UCD1.2 (acc. GCF_002263795.1) and to viral genomes encoding miRNAs that infect cattle, including BLV (acc. NC_001414.1), BFV (acc. NC_001831.1), BoHV-1 (acc. NC_063268.1) and BoHV-5 (acc. NC_005261.3). No mismatches were allowed in the first 18 bases of the read (left end). After alignment to the reference sequences, reads were deduplicated based on their mapping coordinates and the previously extracted UMI tags using UMICollapse v1.0.0 (66).

### miRNAs identification and quantification

Deduplicated reads were mapped to the reference precursor miRNA sequences for bovine, BLV, BFV, BoHV-1 and BoHV-5 obtained from miRBase v22.1 (67). Mature miRNA expression was quantified using miRDeep2 v0.1.3 (68) with default parameters, generating a count matrix of miRNA expression for each sample.

### Differential miRNA Expression analysis

The raw count matrix for all samples was analyzed using DESeq2 v1.44.0 (69) to assess differential miRNA expression between BLV-infected and non-infected animals. Based on miRNA expression profiles, an exploratory analysis of variation patterns between infected and non-infected animals was performed using principal component analysis (PCA). To ensure homoskedasticity of the data, raw miRNA counts were transformed using the Variance Stabilizing Transformation (VST) method implemented in DESeq2 v1.44.0. Pairwise distances between samples were calculated based on the root-mean-square deviation of miRNA expression levels.

To test for differential miRNA expression, the DESeq2 model adjusts miRNA counts by accounting for sample-specific differences, such as library size, sequencing composition, and miRNA-specific biases, using negative binomial models. Under the null hypothesis, the model assumes that a miRNA has the same mean expression across the two conditions (BLV(-) and BLV(+) groups). The statistical significance of the Log_2_ fold change (Log_2_FC) between conditions was assessed using the Wald test. Correction for multiple comparisons was performed using the Benjamini-Hochberg false discovery rate (BH-FDR) method (70). The miRNAs with BH-FDR-adjusted p-values (q) < 0.05 and a Log_2_FC > |1.5| were considered statistically significant. The PCA and Volcano plots were performed under the R environment (71).

### miRNAs gene target prediction

Significant differentially expressed bovine and viral-derived miRNAs between BLV(+) and non-infected cows were used to predict potential bovine gene targets *in silico* (miRNA:mRNA interactions) using miRanda v3.3 (72), PITA v6.0 (73), and RNAhybrid v2.1.1 (74), with default parameters. The analysis utilized 3’-UTR sequences of 21,400 bovine genes from the ARS-UCD1.2 annotation (75). Each gene target prediction algorithm employed a distinct strategy: miRanda relied on seed-site alignment and the thermodynamic stability of RNA-RNA duplexes; PITA incorporated target-site accessibility; and RNAhybrid uses the minimum free energy of hybridization between miRNA-mRNA sequences. Predicted targets were filtered using the following threshold: a score of ≥140 and free energy ≤−20 kcal/mol (miRanda) and free energy ≤−20 kcal/mol (PITA and RNAhybrid). Only gene targets predicted by all three tools (consensus gene targets) were retained for downstream analysis

### Gene ontology and graph-based pathway analysis of miRNA target genes

The functional annotation of the potential target genes was carried out using STRINGdb (76) with Gene Ontology (GO) terms and Kyoto Encyclopedia of Genes and Genomes (KEGG) pathways (77). Next, a protein-protein interaction (PPI) network of the identified target genes was constructed using STRINGdb.

This platform integrates curated and predicted interactions from multiple sources, including experimental evidence, computational predictions, scientific literature, co-expression data and primary public databases. Highly interconnected protein clusters within the network were identified using the Markov Cluster Algorithm (MCL) with default parameters (inflation parameter = 2.5). Only clusters with more than 15 nodes (genes) were considered. GO terms and KEGG pathways overrepresentation analysis was performed on the identified clusters using Fisher’s exact test based on the hypergeometric distribution, using a term similarity threshold > 0.7 (78). Wordclouds plots of significant GO terms grouped by semantic similarity were obtained using the rrvgo package with default parameters (79). A GO term category–gene target graph was generated to visualize the connections between significantly enriched GO terms and their associated genes using the R graphical environment (71).

## Results

In this study, four animals consistently exhibited both high antibody reactivity across all sampling times (IND_5671 = 178.6 ± 28.4; IND_5841 = 191.3 ± 32.8; IND_6021 = 169.8 ± 42.7 and IND_6097 = 183.2 ± 25.4) and high proviral loads at T-3 and T0 (average proviral load: IND_5671 = 63,148.0; IND_5841 = 44,234.3; IND_6021 = 52,902.4 and IND_6097 = 91,255.4). These cows were classified as BLV(+). In contrast, three animals that consistently tested negative for both anti-BLV ELISA and qPCR were classified as BLV(-).

Total RNA quality was confirmed (RIN 7.8–9.6), and small RNA libraries were prepared using UMI-tagged adapters. High-quality, non-redundant reads were obtained through a multi-step processing pipeline. A descriptive characterization of each sample, the number of sequencing reads before and after quality control; merging and deduplication; and mapping statistics are summarized in Table 1.

**Table 1.** Sample characterization and small RNA sequencing output.

| Sample_ID | PR (SD)<sup>1</sup> | PVL | RIN | Raw Reads (n) | Reads after QC, merging and deduplication (n) | Reads mapped to bovine (%) / viral mature miRNAs<sup>2,3</sup> (%) | Total number of identified miRNAs (viral-miRNAs) |
| --- | --- | --- | --- | --- | --- | --- | --- |
| 5671 | 178.6(28.4) | 63,148.0 | 9.6 | 8,090,674 | 4,243,257 | 2,353,523 (96.53) / 81,744 (3.47) | 470 (10) |
| 5841 | 191.3(32.8) | 44,234.3 | 9.6 | 11,869,092 | 4,038,887 | 2,556,676 (97.08) / 74,665 (2.92) | 403 (10) |
| 6021 | 169.8(42.8) | 52,902.4 | 7.8 | 15,487,902 | 6,148,370 | 4,484,871 (92.62) / 331,369 (7.38) | 503 (10) |
| 6097 | 183.2(25.4) | 91,255.4 | 9.3 | 12,511,860 | 4,744,543 | 3,485,727 (94.31) / 198,477 (5.69) | 460 (10) |
| 5830 | <25 | - | 9.0 | 11,903,095 | 5,115,654 | 3,835,811 (99.995) / 215 (0.005) | 503 (0) |
| 6493 | <25 | - | 9.3 | 10,871,552 | 4,312,484 | 3,180,484 (99.994) / 190 (0.006) | 482 (0) |
| 6962 | <25 | - | 9.0 | 11,545,292 | 4,950,884 | 3,575,583 (99.994) / 224 (0.006) | 510 (0) |

<sup>1</sup>Anti-BLV enzyme-linked immunosorbent assay % of reactivity; % of reactivity values < 25.0 indicated negative samples
Abbreviations: ID = sample identification; PR = % of reactivity; SD = standard deviation; RIN = RNA integrity number; PVL = T-3 and T0 BLV average proviral load; QC = quality control.
<sup>2</sup> https://www.mirbase.org/.
<sup>3</sup> A total of 95.7% of reads from BLV(+) samples (n = 18,402,926) and 99.9% of reads from BLV(-) samples (n = 14,378,553) aligned at least once to the Bos taurus reference genome (acc. ARS-UCD1.2). Number of reads aligned to the BLV (acc. NC\_001414.1) and BFV (acc. NC\_001831.1) reference genomes: 648,383 reads from all BLV(+) samples showed at least one alignment to BLV, while only 600 reads from BLV(-) cows mapped to the same genome; regarding the BFV, 138,138 reads from BLV(+) and 141 reads from BLV(-) samples had at least one alignment..

Demultiplexed reads were mapped to the reference miRNA sequences of bovine, BLV, BFV, BoHV-1 and BoHV-5 (n = 1060) to identify and quantify the miRNAs present in each sample. On average, 351 miRNAs (with mapped reads ≥ 5) derived from both bovine and viral sources were identified and quantified across all samples (Supp. Table S1). A multivariate analysis of sample distances based on the normalized miRNA expression matrix revealed that BLV(-) samples clustered together, forming a distinct group separate from BLV(+) samples. PC1 explains 62% of the variance in the expression matrix, while PC2 accounts for 17% (Figure 1).

**Figure 1.**
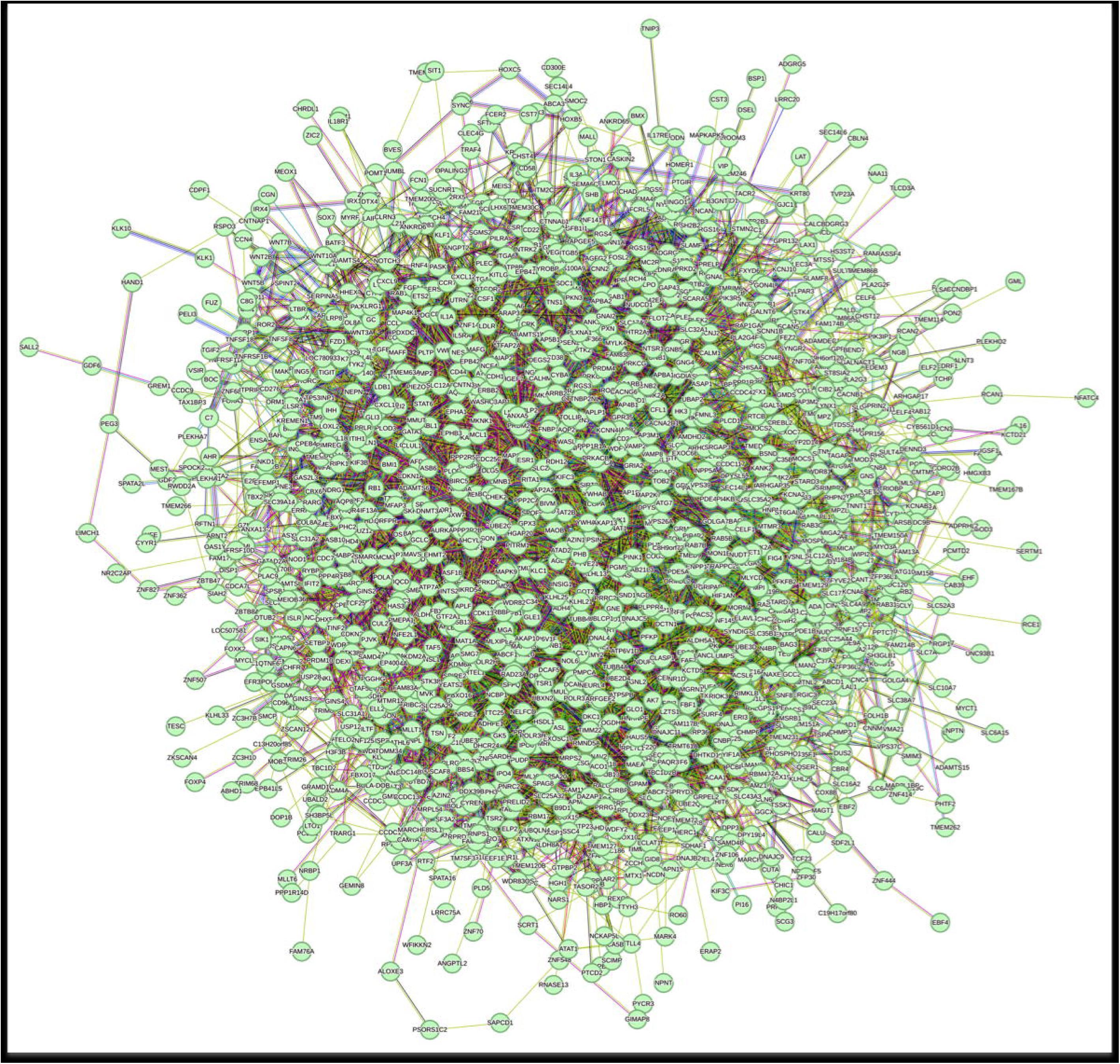
Principal component analysis of miRNAs expression matrix. Blue triangles and Red circles represent BLV(+) and BLV(-) samples, respectively. PC = principal component. PC1 and PC2, represent 62% and 17% of variance contained in the expression matrix data, respectively.

In the BLV(+) group, a total of 10 viral-derived miRNAs were consistently expressed, including 7 from BLV: blv-miR-B1-3p (avgexp = 27,681.1), blv-miR-B2-5p (avgexp = 59,058.0), blv-miR-B2-3p (avgexp = 112.5), blv-miR-B3-5p (avgexp = 309.6), blv-miR-B3-3p (avgexp = 49,312.2), blv-miR-B4-3p (avgexp = 65.1), and blv-miR-B5-5p (avgexp = 951.7). Additionally, 3 BFV-derived miRNAs were identified: *bfv-* miR-BF1-5p (avgexp = 6,844.7), bfv-miR-BF1-3p (avgexp = 8,257.3), and bfv-miR-BF2-5p (avgexp = 14,920.6). In contrast, viral-derived miRNAs expression was low (avgexp < 30.0) or undetectable in the BLV(-) group, except for blv-miR-B2-5p (avgexp = 67.4) and blv-miR-B3-3p (avgexp = 49.8) (Table 2).

**Table 2.** Viral derived-miRNAs expression.

| Condition | BLV(+) |  |  |  |  | BLV(-) |  |  |  |
| --- | --- | --- | --- | --- | --- | --- | --- | --- | --- |
| miRNA/ID | 5671 | 5841 | 6021 | 6097 | avgexp(SD) | 5830 | 6493 | 6962 | avgexp(SD) |
| bfv-miR-BF1-5p | 8,973.2 | 719.9 | 14,813.3 | 2,872.6 | 6,844.7(6,359.3) | 15.6 | 5.8 | 15.4 | 12.3(5.6) |
| bfv-miR-BF1-3p | 12,947.2 | 1,329.6 | 15,826.4 | 2,925.9 | 8,257.3(7,204.2) | 7.1 | 11.6 | 8.6 | 9.1(2.3) |
| bfv-miR-BF2-5p | 17,694.3 | 1,735.7 | 33,383.7 | 6,868.5 | 14,920.6(13,991.2) | 0.0 | 15.4 | 17.1 | 10.8(9.4) |
| bfv-miR-BF2-3p | 2.8 | 0.0 | 6.3 | 6.2 | 3.8(3.0) | 0.0 | 0.0 | 0.0 | 0.0(0.0) |
| blv-miR-B1-5p | 0.0 | 0.0 | 0.0 | 0.0 | 0(0.0) | 0.0 | 0.0 | 0.0 | 0.0(0.0) |
| blv-miR-B1-3p | 7,556.4 | 13,726.1 | 46,243.5 | 43,198.3 | 27,681.1(19,875.4) | 28.3 | 24.1 | 24.0 | 25.5(2.5) |
| blv-miR-B2-5p | 30,941.9 | 35,856.1 | 88,113.4 | 81,320.4 | 59,057.9(29,825.4) | 60.9 | 69.4 | 71.9 | 67.4(5.7) |
| blv-miR-B2-3p | 34.2 | 47.4 | 171.3 | 197.0 | 112.5(83.6) | 0.0 | 0.0 | 0.0 | 0.0(0.0) |
| blv-miR-B3-5p | 262.0 | 371.8 | 259.8 | 344.7 | 309.6(57.3) | 0.0 | 0.0 | 0.0 | 0.0(0.0) |
| blv-miR-B3-3p | 37,074.5 | 33,573.4 | 61,540.1 | 65,060.6 | 49,312.2(16,278.9) | 39.7 | 55.9 | 53.9 | 49.8(8.8) |
| blv-miR-B4-5p | 1.4 | 0.0 | 0.0 | 1.0 | 0.6(0.7) | 0.0 | 0.0 | 0.0 | 0.0(0.0) |
| blv-miR-B4-3p | 39.9 | 69.9 | 79.8 | 70.8 | 65.1(17.4) | 0.0 | 0.0 | 0.0 | 0.0(0.0) |
| blv-miR-B5-5p | 864.3 | 972.1 | 1,218.4 | 752.0 | 951.7(199.2) | 0.7 | 1.0 | 0.9 | 0.8(0.1) |
| blv-miR-B5-3p | 0.0 | 1.2 | 0.8 | 2.1 | 1.0(0.9) | 0.0 | 0.0 | 0.0 | 0.0(0.0) |

Abbreviations: ID = Sample ID; avgexp = normalized mean expression; SD = Standard deviation

The mapping of all reads from BLV(-) samples to the bovine reference genome (acc. ARS-UCD1.2) showed that only 716 out of 14,378,553 reads failed to align, indicating that nearly all reads were derived from bovine small RNAs. Besides, the number of reads aligned to the BLV (acc. NC_001414.1) and BFV (acc. NC_001831.1) reference genomes from BLV(-) cows were only 600 and 141 reads, respectively (Table 1). Moreover, the alignment of the same reads dataset to the identified BLV- and BFV-miRNAs resulted in 642 reads with at least one aligment. Among these, only six reads also map to the bovine reference genome, and none aligned any other known miRNA from any organism in the miRBase database (n = 48,871 mature miRNAs). The Supp. Figure S2 shows the coverage of all reads from BLV(+) and BLV(-) cows across the reference genomes of BLV and BFV, emphasizing that detectable read coverage is restricted to known miRNA regions for both viral genomes.

Differential expression analysis of host miRNAs, identified a total of seven bovine miRNAs (bta) with a fold change (FC) > |1.5| and a q-value < 0.05 as differentially expressed (BTA-miRNAs-DE) between the BLV(+) and BLV(-) groups: bta-miR-375, bta-miR-133a, bta-miR-677, bta-miR-1, bta-miR-3613a, bta-miR-9-5p, and bta-miR-95 (Figure 2; Supp. Table S2). In Supp. Figure S3, a heatmap was generated to visualize the expression of miRNAs in BLV(+) and BLV(-) cows. A hierarchical clustering was applied to both dimensions to show groups of samples with similar expression profiles, as well as clusters of miRNAs with coordinated expression patterns.

**Figure 2.**
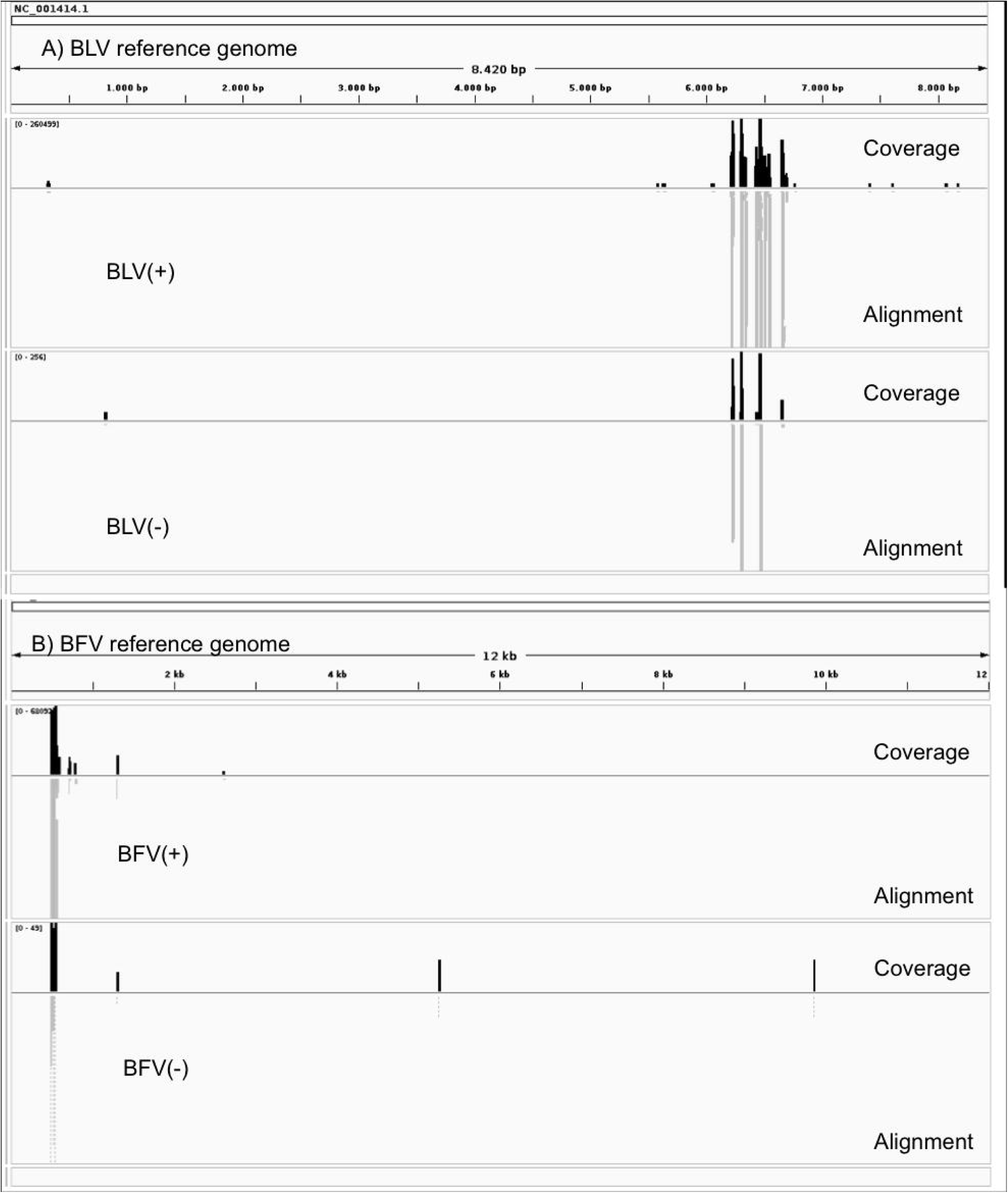
Volcano plot summarizing the results of the differential expression analysis of bovine miRNAs between BLV(+) and BLV(-) groups. Each circle represents a bovine miRNA. Red circles indicate miRNAs with FC < |1.5| and non-significant, while blue circles represent miRNAs with FC > |1.5| and q < 0.05 (-log(q-value) > 1.3). Viral-derived miRNAs, primarily quantified in BLV(+) samples, were excluded from the plot. Vertical dashed lines represent a |1.5| FC in miRNA expression. Horizontal dashed line represents the significance cutoff q-value = 0.05.

To evaluate the potential functional impact of the seven differentially expressed BTA-miRNAs and the ten viral-derived miRNAs, a target gene prediction analysis was performed. A total of 1,518 genes were identified as putative targets, based on the consensus of three independent prediction tools. Among these, 281, 977, and 260 genes were exclusively targeted by BTA-miRNAs-DE, BLV-miRNAs, and BFV-miRNAs, respectively (Supp. Table S3). No overlap was observed among the predicted target. Then, functional annotation of the 1,518 potential target genes was conducted using Gene Ontology (GO) terms and metabolic pathways, revealing 91 significantly overrepresented GO biological process (BP) terms, 10 GO molecular function (MF) terms and 30 GO cellular component (CC) terms. The word cloud visualizations for the enriched GO terms in the GO BP and GO MF categories are shown in Figure 3, the most frequent occurring words for the overrepresented processes and functions included “Regulation”, “Process”, “Response”, “Binding”, “Activity”, among others. The overrepresentation analysis of KEGG and Reactome metabolic pathways revealed four significant terms: “*Ras signaling pathway*” (bta04014, q = 0.04), “*Pathways in cancer*” (bta05200, q = 0.02), “*Oxytocin signaling pathway*” (bta04921, q = 0.024), and “*Cushing syndrome*” (bta04934, q = 0.04). Additionally, the terms “*Signal Transduction*” (BTA-162582, q = 0.0002) and “*Metabolism*” (BTA-1430728, q = 0.01) were significant in Reactome pathways analysis (Supp. Table S4).

**Figure 3.**
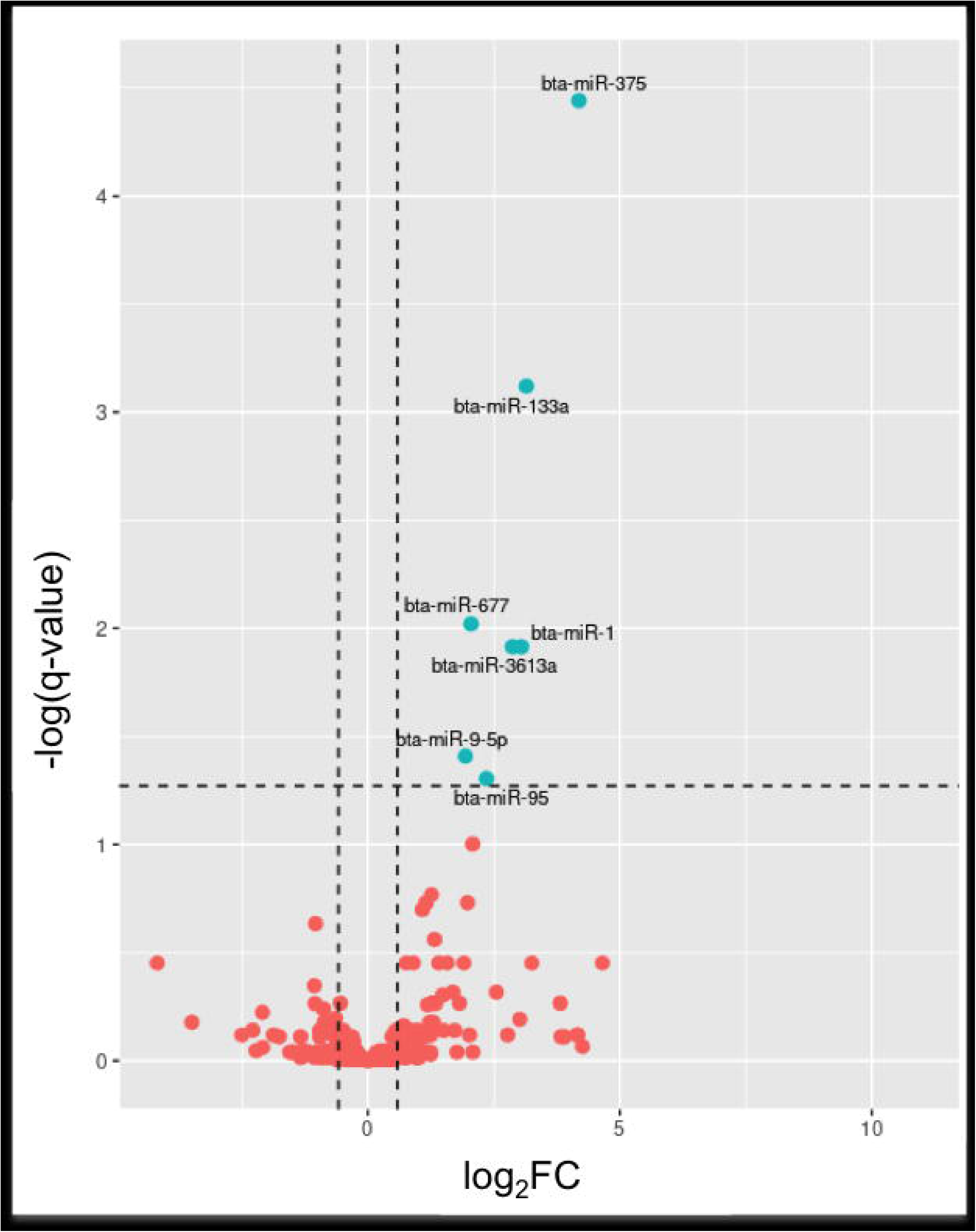
Word cloud for GO term overrepresented in target genes for Biological Process A) and Molecular Fuction B) categories.

A protein-protein interaction (PPI) network was developed, comprising 1,518 nodes (genes) and 6,075 edges (interactions) (Supp. Figure S2). From this network, three highly interconnected protein clusters (C1, C2, and C3) were identified (PPI p-value < 1.0e^-16^), which potentially represent hubs of specialized biological function (Figure 4). Functional overrepresentation analysis for these clusters revealed key cellular processes involved in immune response modulation, cell signaling, and mechanisms related to cancer development (Supp. Table S4). Interestingly, despite the lack of overlap among target genes of BTA-, BLV-, and BFV-derived miRNAs, their predicted functions converge on interconnected hubs, indicating potential interplay at the pathway level. The enrichment graph plot depicts the interconnections among the top 15 enriched BP GO terms per cluster, highlighting functional overlap through target genes (Figure 5).

**Figure 4.**
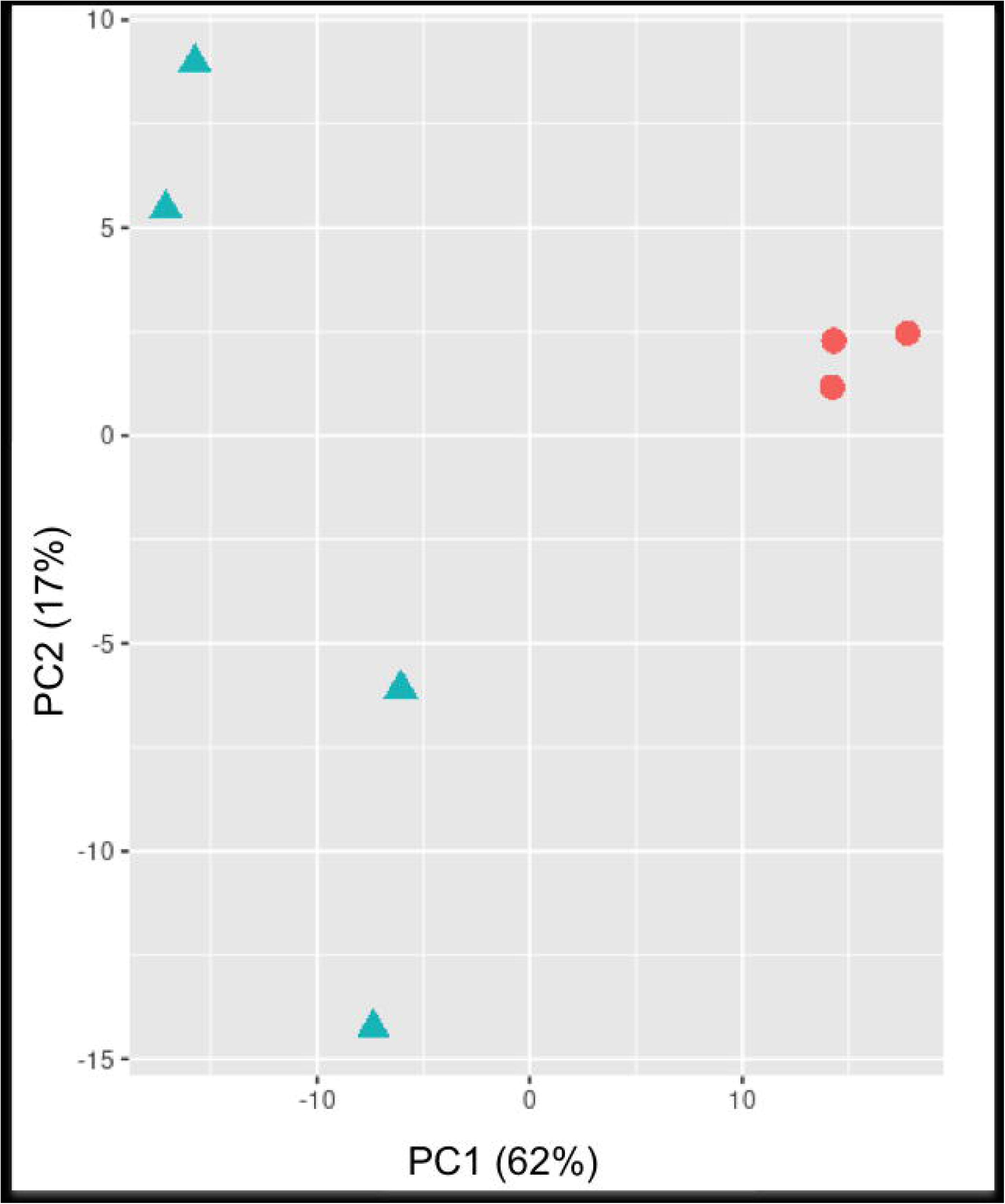
Protein-protein interaction (PPI) cluster analysis. Three cluster (> 15 nodes) were identified A) Cluster 1 (C1): N° nodes: 21 - N° edges: 81, avg. node degree: 7.7; B) Cluster 2 (C2): N° nodes: 20 – N° edges: 64, avg. node degree: 6.4; C) Cluster 3 (C3): N° nodes: 16 – N° edges: 34, avg. node degree: 4.2. Green nodes: predicted BTA-miRNA-DE protein target; Blue nodes: predicted BFV-miRNA protein target; Red nodes: predicted BLV-miRNA protein target. Edges represent protein-protein interactions as determined by STRINGdb evidence, including: known interactions (curated databases and/or experimentally validated), predicted interactions (gene fusions, gene co-occurrence), text mining, co-expression, and protein homology.

**Figure 5.**
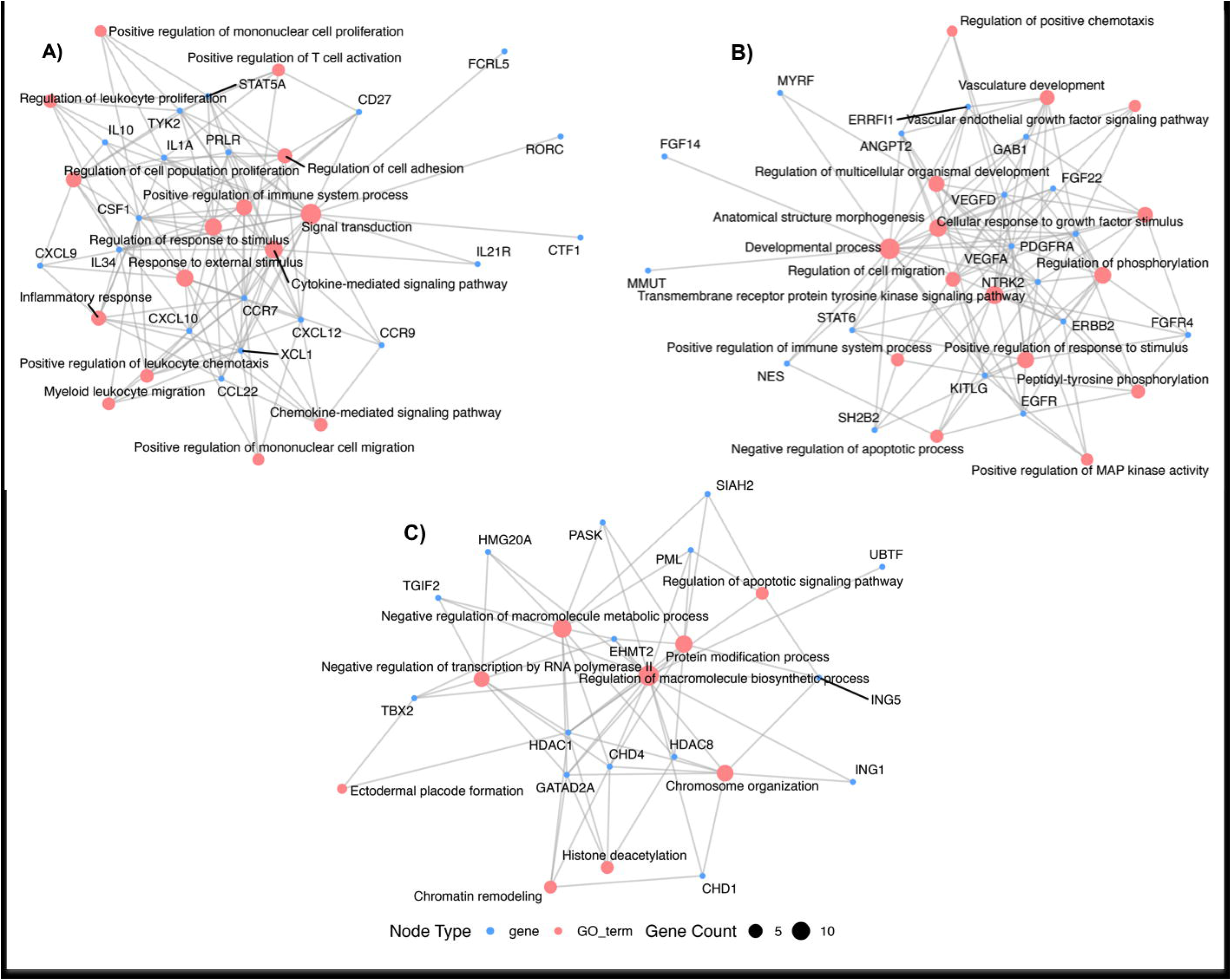
Enrichment graph plot of PPI clusters. Each graph shows the relationship between the 15 top enriched BP GO terms (light pink node) and their associated gene (light blue nodes). Node size reflects the number of genes associated with each GO term (Count), and edges indicate gene-GO term associations. Genes in each graph include predicted gene targets of A) BTA-miRNA-DE, B) BLV-miRNA, and C) BFV-miRNA, respectively.

For example, significantly overrepresented GO BP terms for C1 (q < 0.05), included “*Cytokine-mediated signaling pathway*”, “*Signal transduction*”, “*Positive regulation of leukocyte chemotaxis*”, “*Positive regulation of immune system process*”, “*Regulation of leukocyte proliferation*”, “*Regulation of cell adhesion*”, “*Myeloid leukocyte migration*”, “*Positive regulation of T cell activation*”, “*Regulation of leukocyte apoptotic process*”, “*Regulation of interferon-gamma production*”, amog others, focusing especially on processes related to immune activation, cytokine production, and immune regulation. For C2, key terms represent biological processes primarily tied to cell growth, development, and tissue morphogenesis, such as “*Transmembrane receptor protein tyrosine kinase signaling pathway*”, “*Vascular endothelial growth factor signaling pathway*”, “*ERBB2-EGFR signaling pathway*”, “*Positive regulation of MAP kinase activity*”, “*Hemopoiesis*”, “*Negative regulation of apoptotic process*”, “*Anatomical structure morphogenesis*”, “*Positive regulation of epithelial cell proliferation*”, “*Positive regulation of mast cell chemotaxis*”, among others.

In C3, the terms are mostly associated with epigenetic control and transcriptional regulation through chromatin modification, including “*Histone deacetylation*”, “*Histone modification*”, “*Chromatin remodeling*”, “*Negative regulation of transcription by RNA polymerase II*”, “*Chromosome organization*”, ”*Regulation of apoptotic signaling pathway*”, among others.

Significantly enriched KEGG and Reactome pathways for clusters included, C1: “*Cytokine-cytokine receptor interaction*” (bta04060, q-value = 8.40e^-22^), “*Viral protein interaction with cytokine and cytokine receptor*” (bta04061, q-value = 1.02e^-16^), “*Cytokine Signaling in Immune system*” (BTA-1280215, q-value = 1.09e^-06^), “*Chemokine receptors bind chemokines*” (BTA-380108, q-value = 1.10e^-06^), “*Interleukin-21 signaling*” (BTA-9020958 q-value = 0.0057), “*IL-6-type cytokine receptor ligand interactions*” (BTA-6788467, q-value = 0.0105); C2: “*MAPK signaling pathway*” (bta0401, q-value = 3.50e^-10^), “*Regulation of actin cytoskeleton*” (bta04810, q-value = 0.0014), “*HIF-1 signaling pathway*” (bta04066, q-value = 0.0029), “*Signaling by Receptor Tyrosine Kinases*” (BTA-9006934, q-value = 3.13e^-16^), “*PI5P, PP2A and IER3 Regulate PI3K/AKT Signaling*” (BTA-6811558, q-value = 1.62e^-09^), “*PI3K Cascade*” (BTA-109704, q-value = 0.0034), “*Regulation of KIT signaling*” (BTA-1433559, q-value = 0.0058); and C3: “*Regulation of TP53 Activity through Acetylation*” (BTA-6804758, q-value = 2.85e^-08^), “*RNA Polymerase I Transcription Initiation*” (BTA-73762, q-value = 1.02e^-07^), “*Gene expression (Transcription)*” (BTA-74160, q-value = 3.05e^-05^), “*Chromatin modifying enzymes*” (BTA-3247509, q-value = 4.15e-^05^), “*Regulation of PTEN gene transcription*” (BTA-8943724, q-value = 0.00052).

## Discussion

In this study, we performed a comprehensive characterization of circulating miRNAs in the peripheral blood of naturally BLV-infected and non-infected cattle. Notably, all BLV-infected cows included in the analysis consistently exhibited HPVL levels across time points; a condition previously associated with BLV pathogenesis progression (80; 81) and increased risk of viral transmission within herds (82). Following the identification of BLV-miRNAs in persistently infected cell lines (21), their expression was assessed using next-generation sequencing for complete miRNA profiling and RT-qPCR to test the expression of candidate miRNAs. BLV-miRNAs were identified in primary leukemic B-cells and B-cell lymphomas isolated from BLV-infected ovine/bovine tumors (44), experimentally infected cattle (83), and naturally BLV-infected cattle (45; 84; 85).

On the other hand, BFV-miRNAs have been identified in both persistently and recently infected MDBK cells, as well as in BFV experimentally challenged cattle (23). However, to our knowledge, this is the first report of the co-expression of miRNAs derived from two distinct viruses (BLV and BFV) in naturally infected cattle. We identified and quantified seven BLV-derived and three BFV-derived miRNAs. No miRNAs from BoHV-1 and −5 were detected.

Coinfections of BLV and BFV are prevalent among cattle (53), akin to the occurrences in cats infected with Feline foamy virus (FFV) and Feline leukemia virus (FeLV) (86), as well as in baboons with Simian foamy virus (SFV) and Simian T-cell leukemia virus (STLV) (87). While BFV infections in cattle are generally regarded as mild and asymptomatic (23; 53), there is evidence to suggest that co-infections with BFV and BLV could amplify pathogenicity and impair the bovine immune system, thereby aiding in the transmission and spread of BFV (53; 88; 89; 90). Infectious BFV has been extracted from raw milk, and considering that foamy viruses can cross species barriers, there is rising concern over the zoonotic potential of BFV in humans (91; 92). Moreover, increased pathogenic effects have been noted in mixed infections of macaques with SFV and Simian immunodeficiency virus (SIV) (93), as well as in cats coinfected with Feline immunodeficiency Virus (FIV) and FFV or FeLV and FFV (86; 94).

The seven miRNAs derived from BLV comprised no more than 8% of the total miRNA sequencing reads in all analyzed samples (data not provided), consistent with the expression levels reported by Casas et al. (2020) (85) for natural BLV infections. In contrast, Rosewick et al. (2013) (44) found that all ten mature BLV-derived miRNAs accounted for about 40% of total miRNAs present in B-cell lymphomas from sheep infected with BLV. Similarly, Ochiai et al. (2021) (95) noted that BLV-derived miRNAs made up 38% of total miRNAs in Japanese black cattle diagnosed with enzootic bovine leukosis (B-cell lymphoma). Additionally, Kincaid et al. (2012) (21) reported that BLV miRNAs were found in greater quantities than cellular miRNAs in the BL3.1 cell line persistently infected with BLV. Although the relative expression levels of individual BLV miRNAs differ across studies, blv-miR-B4-5p and blv-miR-B5-3p appear to be either weakly expressed or undetectable. The dominant BLV-miRNA may change depending on the disease stage or phase of infection analyzed. Overall, these findings suggest that BLV miRNAs are highly expressed in *ex vivo* cell cultures, persistently infected cell lines, primary B-cell tumors, and cattle affected by enzootic bovine leukosis. In cattle with subclinically BLV-infections, their expression levels might vary based on several factors, including the infection route (natural or experimental) and the animal’s age (whether calf or adult cow), among other factors (21; 44; 83; 84; 85; 95). Conversely, the other BLV transcripts show little to no expression *in vivo*, highlighting the essential function of BLV miRNAs in viral persistence (96; 97).

Likewise, three BFV-derived miRNAs (bfv-miR-BF1-5p, bfv-miR-BF1-3p, and bfv-miR-BF2-5p) have been identified in MDBK cells that are persistently and recently infected with BFV, showing higher expression levels during persistent infections. Furthermore, two BFV-miRNAs were found in the peripheral blood leukocytes of cattle experimentally infected with BFV, utilizing a qPCR assay (23). This research is the first to document the expression of these three BFV-derived miRNAs in natural setting infections.

A previous study compared BLV seropositive and seronegative cattle but without testing for BLV genomic material (85). The detection of BLV-miRNAs in seronegative cows suggests a previous exposure to the virus, likely resulting in an inadequate immune response that did not generate a positive result in the anti-BLV ELISA test. However, a longitudinal analysis of seroconversion was not performed. This study found that all ten virus-derived miRNAs were consistently and highly expressed in the BLV(+) group, whereas only a few showed weak expression in the BLV(-) group. Although undetected infections cannot be entirely excluded, it is improbable that adult animals over two years old, which tested negative via qPCR and ELISA at two intervals (three months apart), were unrecognized cases. The near-complete alignment of BLV(-) reads to the bovine genome confirms their bovine origin, while a small subset of reads mapped exclusively to viral miRNAs. Therefore, it suggests that the reads aligning to BLV- and BFV-derived miRNAs in BLV(-) samples are likely bona fide and not the result of cross-mapping or artifacts (data not shown). Moreover, sample mislabeling appears to be unlikely, if such an issue had occurred, it would expect to observe expression profiles of viral miRNAs in BLV(+) samples that mirrored those of BLV(-). However, our data did not show such patterns. The miRNA expression profiles in BLV(-) samples were not consistent with those from BLV(+) samples, further arguing against sample misidentification or cross-contamination. The potential for BLV-or BFV-miRNAs to function as xeno-miRs and be transmitted from infected to uninfected animals remains a question for future research.

Notably, the expression profiles of BLV(+) / BFV-miRNAs(+) and BLV(-) samples distinctly differentiate the two groups of cows groups, as shown in the PCA plot. This indicates a shift in the host miRNA expression profile between them.

Among the significant differentially expressed bovine miRNAs, bta-miR-375 was identified, which has been previously reported to be associated with BLV infection in cattle (85; 95; 98), and suggested it as an early biomarker for diagnosing enzootic bovine leukosis (EBL). Moreover, its levels effectively distinguished EBL-affected cattle from asymptomatic cattle with high sensitivity and specificity (99).

In a similar manner, bta-miR-133a was found to be differentially expressed in the serum of BLV-seropositive cows compared to seronegative cows (98). The other significantly differentially expressed miRNAs (bta-miR-677, bta-miR-95, bta-miR-9-5p, bta-miR-3613a, and bta-miR-1) have not been previously reported in relation to BLV and/or BFV infections.

To date, efforts to assign biological functions to BLV- and BFV-derived miRNAs have focused on predicting potential target genes for specific BLV- and BFV-miRNAs, in addition to selecting candidate target genes for functional assays (21; 41; 44; 45; 83; 84; 85). In this study, we evaluated the functional interaction (protein-protein interaction network) of potential target genes of BLV-miRNAs, while also considering target genes of BFV-miRNAs and differentially expressed bovine-miRNAs.

It has been demonstrated that BLV-miRNAs are crucial in promoting disease progression in cattle (83; 100). Additionally, BLV fails to induce leukemia/lymphoma in sheeps (oncogenicity suppressed) challenged with an isogenic BLV provirus lacking the miRNA genomic region.

The functional annotation of the proteins within hub C1 revealed that immune system-related GO BP terms, such as “*Cytokine-mediated signaling pathway*”, “*Positive regulation of leukocyte chemotaxis*”, “*Regulation of leukocyte proliferation*”, “*Positive regulation of T cell activation*”, “*Regulation of leukocyte apoptotic process*”, “*Regulation of interferon-gamma production*”, were significant overrepresented, along with the KEGG and Reactome terms: “*Cytokine-cytokine receptor interaction*”, “*Viral protein interaction with cytokine and cytokine receptor*”, “*Cytokine signaling in immune system*”. The Figure 5 A) showed the enrichment graph connecting gene target to BP GO terms. It is widely accepted that BLV infection changes the cytokines expression patterns and alters how the immune system produces cytokines in response to BLV antigen stimulation (101; 102). Particularly, BLV disease progression would be related to the suppression of the cell-mediated immune response (103). For example, Interleukin-10 (IL-10), a suppressor and anti-inflammatory immune response cytokine, is over expressed in cows with persistent lymphocytosis (104), which can inhibit cytokine production by Th1 cells (for example, IL-2, IL-12 and gamma interferon) (104; 105) influencing the B-cell proliferation and differentiation (104; 106).

In turn, the analysis of overrepresented metabolic pathways for the complete potential target gene set identified “*Pathway in cancer*” as one of the significant KEGG pathways. This pathway involves a series of cellular signaling pathways that activate crucial hallmarks of tumorigenesis, including tissue invasion and metastasis, evading apoptosis, genomic instability, cell proliferation, genomic damage, and insensitivity to anti-growth signals (107). The overrepresented GO BP terms for the biological function hub C2 included terms associated with cellular processes that, when disrupted, could be linked to tumor development events. These terms included “*Transmembrane receptor protein tyrosine kinase signaling pathway*”, “*Vascular endothelial growth factor signaling pathway*”, “*ERBB2-EGFR signaling pathway*”, “*Positive regulation of MAP kinase activity*”, “*Hemopoiesis*”, “*Negative regulation of apoptotic process*”, among others. Interestingly, other overrepresented KEGG terms in C2 included the “*MAPK signaling pathway*”, “*JAK-STAT signaling pathway*”, “*HIF-1 signaling pathway*”, “*Regulation of actin cytoskeleton*” and Reactome terms such as “*PI3K/AKT Signaling*” and “*Signaling by Receptor Tyrosine Kinases*”. This signaling pathways are amplified and propagated intracellularly by various kinases, ultimately affecting how transcription factors and histone-modifying complexes control downstream gene expression (108). In this context, a crucial signaling cascade most altered in human cancers is the mitogen-activated protein kinase (MAPK) pathway, which includes the RAS–RAF–MAPK kinase (MEK)–extracellular signal-regulated kinase (ERK) pathway (109; 110). Using the ovine BLV pathogenesis model, sheeps were experimentally challenged with an isogenic BLV-miRNA deletion mutant, and global transcriptome analysis revealed that BLV-miRNAs primarily promote the proliferation of BLV-infected B-cells by inhibiting immune response and cell signaling pathways (100). In turn, in a reporter assay, the FBJ murine osteosarcoma viral oncogene homolog (*FOS*) was identified as a direct target of BLV-miRNAs. The *FOS* gene is a component of the activator protein-1 (AP-1) complex, which is involved in the primary response to B-cell receptor signaling and is frequently downregulated in many cancers, including breast carcinomas (83; 111).

Additionally, enriched GO BP terms across C3 included “*Histone modification*”, “*Chromatin remodeling*”, and “*Histone deacetylation*”, “*Negative regulation of transcription by RNA polymerase II*”, ”*Regulation of apoptotic signaling pathway*”, among others; as well as the Reactome terms “*Regulation of TP53 Activity through Acetylation*” and “*Chromatin modifying enzymes*”. The transcription factor *TP53*, a tumor suppressor activated by DNA damage, plays a critical role in maintaining genomic integrity (112). The expression of two transcription factors, B-lymphocyte-induced maturation protein 1 (*BLIMP1*) and B-cell lymphoma 6 (*BCL6*), is negatively correlated with BLV-miRNA expression in BLV(+) cows. These transcription factors play a pivotal role regulating B-cell differentiation, antibody affinity and T-cells immune function. Particularly, an important function of *BCL6* is to enable GC B-cells to proliferate in response to T-cell antigens by specifically repressing gene expression related to DNA damage sensing. This repression allows for the tolerance of DNA breaks induced during immunoglobulin class rearrangement and somatic hypermutation (84; 113; 114).

The functional role of BFV-miRNAs remains poorly investigated. Predicted gene targets for bfv-miR-BF2-5p were Ankyrin repeat domain-containing protein (*ANKRD17*) and Bax-interacting factor 1 (*BIF1*) (41). *BIF1* plays a key role in activating the pro-apoptotic Bax protein in the intrinsic apoptosis pathway as well as in autophagy and autophagosome formation, thus acting as a tumor suppressor (115; 116). *ANKRD17,* on the other hand, is involved in DNA replication and cell cycle progression. It also interacts with genes that are responsible for sensing viral RNAs and trigger immune responses (117; 118).

Notably, although target genes of bovine, BLV, and BFV miRNAs do not overlap, functional analysis revealed convergence on similar cellular processes, implying coordinated regulatory interactions (Figure 5).

Regarding the differentially expressed BTA-miRNAs, miR-1, miR-133a, and miR-375 are recognized as tumor suppressors and are downregulated in various types of cancer, exerting their effects through distinct mechanisms (119; 120). For instance, miR-1 interacts with the proto-oncogene B-cell lymphoma 2 (*BCL-2*), an anti-apoptotic gene. Overexpression of miR-1 inhibits cell proliferation, migration, and invasion while promoting apoptosis in breast cancer (121). Similarly, miR-133a modulates biological processes such as proliferation, apoptosis, and autophagy (120). On the other hand, miR-375 is significantly downregulated in several cancers and considered a biomarker of poor prognosis (122; 123; 124; 125), but it is overexpressed in breast cancer (126). Similarly, miR-95-3p and miR-9-3p act as prognostic markers, promoting the progression of cervical and breast cancers, respectively (127; 128).

Conversely, the differentially expressed bovine miRNA bta-miR-677 enhances the production of type I interferons (IFN-I) and interferon-stimulated genes (ISGs) (129). Similarly, inhibiting miR-3613-3p decreases the expression of IFN-α and IFN-β, thereby affecting the anti-hepatitis B activity of interferons (130). Moreover, miR-3613-3p has been recognized as a tumor suppressor, with its deletion associated with poor prognosis in estrogen receptor-positive breast cancer patients (131).

## Conclusion

In this study, we report for the first time the co-expression of seven miRNAs derived from BLV and three from BFV in cattle naturally infected with BLV. Several bovine BTA-miRNAs were found to be differentially expressed (DE) between cows with BLV high proviral load (PVL) and non-infected cows. Among them, bta-miR-375 was previously identified as a potential early biomarker for the progression of enzootic bovine leukosis, and this study further emphasizes its potential utility in identifying high PVL cows alongside other candidate miRNAs.

The functional analysis of protein-protein interaction networks involving BLV-, BFV-, and BTA-DE miRNAs-targeted genes identified key metabolic pathways that potentially underlie tumorigenesis and immune response modulation, including cellular signaling pathways, regulation of T cell activation, apoptosis, chromatin remodeling, and cytoskeleton regulation. Moreover, miRNA targets from both host and viral origins would act synergistically to influence cellular processes. The co-expression of miRNAs derived from BLV and BFV raises concerns about whether their interaction may exacerbate disease pathogenesis or facilitate the dissemination of BFV in light of the zoonotic potential that BFV presents. Ultimately, the identified miRNAs and associated metabolic pathways offer promising opportunities for developing diagnostic tools and therapeutic strategies aimed at controlling viral persistence and tumorigenesis in cattle.

## Declarations

### Competing interests

The authors declare that they have no competing interests.

### Funding

The present study was partially supported by the Instituto Nacional de Tecnología Agropecuaria (INTA, Argentina) projects 2023-2017-PD-I113 and 2023-2027-PD-I114, the Fondo para la Investigación Científica y Tecnológica (FONCyT) PICT-2017-0262, and by the Consejo Nacional de Investigaciones Científicas y Técnicas (CONICET, Argentina) PIP 2021-2023-11220200101106CO01

### Authors’ contributions

Conceptualization: DDG, CM, JPJ, HAC.

Data curation: HAC. Formal analysis: MIP, HAC.

Funding acquisition: CM, JPJ, HAC

Investigation: MIP, GSA, CP, DDG, KGT, CM, JPJ, HAC.

Methodology: HAC. Visualization: HAC.

Writing – original draft: HAC.

Writing – review & editing: MIP, GSA, CP, DDG, KGT, CM, JPJ, HAC. Roles as defined by: CRediT (contributor role taxonomy)

### Availability of data

Raw small sequencing reads are available in the NCBI Sequence Read Archive (SRA) under BioProject accession number PRJNA1219522.

## Acknowledgments

We would like to thank Dr. Andrea Puebla and Pablo Vera from the Genomic Unit (IABIMO, INTA-CONICET) for their technical assistance with the small RNA next-generation sequencing assay.

---

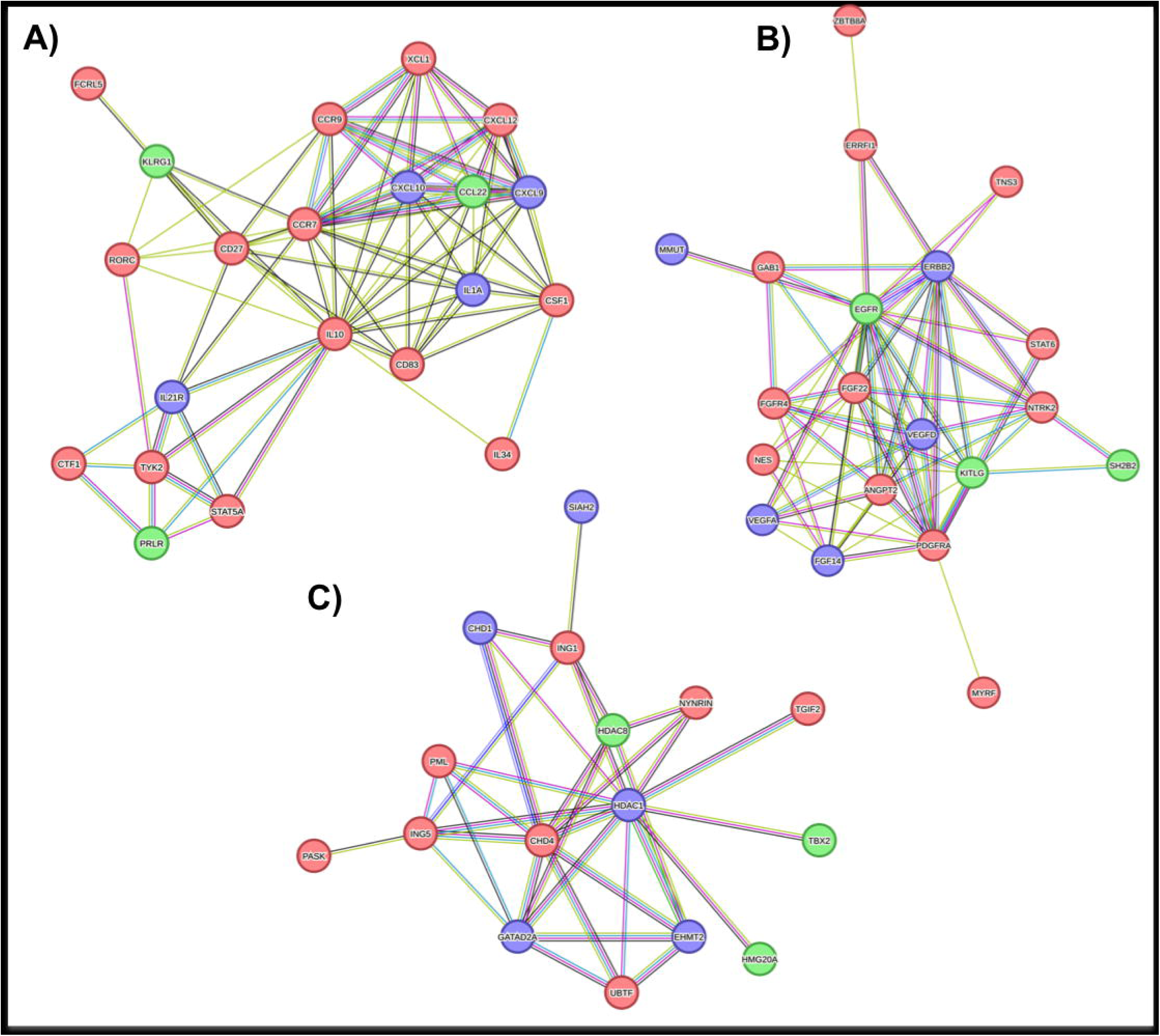

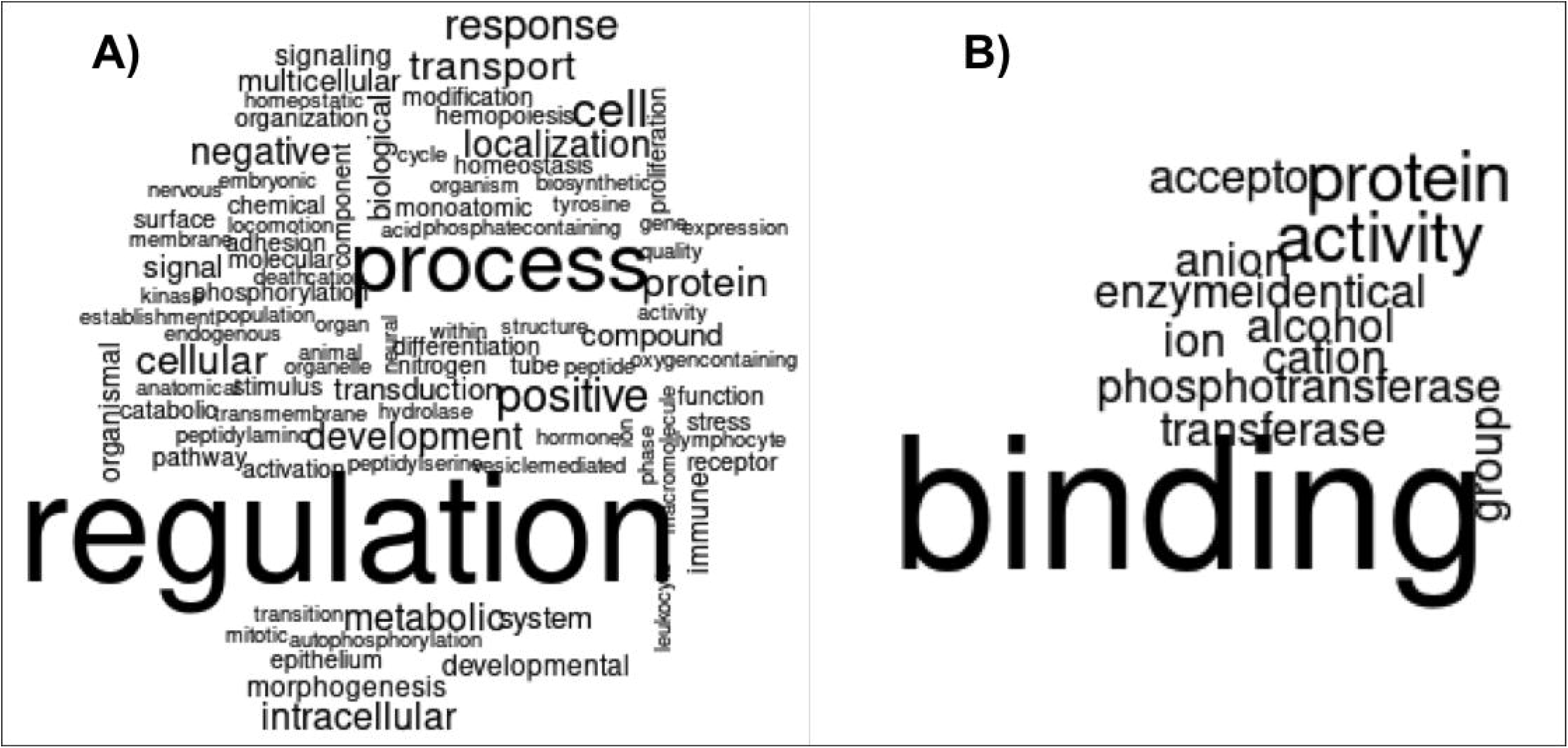

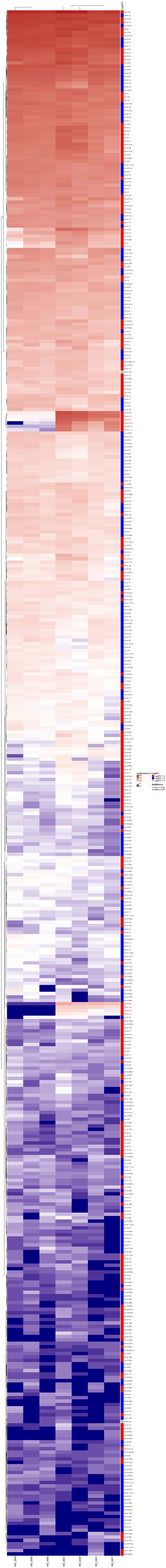

## Notes

### Competing Interest Statement

The authors have declared no competing interest.

### Summary of Updates

We have made several revisions to the manuscript; among them, the following were the main changes: Clarification of animal selection criteria for small RNA sequencing We revised the Methods section to include a new paragraph (L131,144) detailing how animals were screened and selected. This addition aims to improve transparency regarding the selection criteria and to better support the validity of our experimental design. Refinement of miRNA target prediction to reduce potential biases To rule out potential biases or artifacts in the target prediction process, we updated our analysis to use three widely adopted and complementary miRNA target prediction tools, together with strict filtering criteria. Only targets predicted by all three tools (consensus targets) were retained for downstream analysis, reducing the likelihood of tool-specific false positives (Methods section, L232,241). Assessment of viral miRNA expression in BLV(-) cows We further examined the expression of viral miRNAs in BLV(-) cows by analyzing unmapped reads. Of the 14,378,553 reads from BLV(-) samples, only 716 did not align to the bovine reference genome. When these were aligned to the BLV and BFV miRNAs, only 642 reads were mapped, of which just six also aligned to the bovine genome and none matched other mature miRNA sequences in the miRBase database (48,871 entries). While mapping artifacts can never be completely ruled out, the extremely low number and specificity of these reads suggest such occurrences are rare and unlikely to impact our conclusions (L283,293). We have also added Supplementary Figure S2 showing that read coverage from BLV(+) and BLV(-) cows is limited to known miRNA regions in the BLV and BFV genomes.

